# Investigation of CNS damage following HIV infection and methamphetamine exposure using human iPSC-derived microglia and 3D cerebral assembloid models

**DOI:** 10.64898/2025.12.06.692767

**Authors:** Shashi Kant Tiwari, Gulshanbir Baidwan, Samantha Trescott, Shweta Jakhmola, Nicole G. Coufal, Tariq M. Rana

## Abstract

HIV-associated neurocognitive disorder (HAND) is characterized by glial activation and neuroinflammation emerging from HIV infection. Moreover, methamphetamine (METH) is a highly addictive psychostimulant whose use is linked to high HIV prevalence, significantly worsening clinical outcomes for people living with HIV and hastening onset of systemic illness. Overall, pathologies associated with HIV infection/METH use comorbidity include neuroinflammation, neurotoxicity, synaptodendritic damage, and resulting cognitive dysfunction. However, mechanisms underlying this neuropathogenesis remain elusive due to the scarcity of human brain-specific experimental model systems. Therefore, we created a next-generation three-dimensional (3D) human brain model derived from human induced pluripotent stem cells (hiPSCs). Specifically, this 3D *in vitro* cerebral assembloid model integrates functional microglia derived from hiPSCs with cerebral organoids. Microglia are key contributors to HAND symptoms associated with HIV/METH comorbidity. We show that the presence of microglia in this assembloid model allows productive infection with HIV, which is enhanced by METH exposure, resulting in increased glial activation, inflammatory responses (namely IL-1β and IL-6 release), and neurotoxicity marked by neuronal cell death and synaptic protein loss. Importantly, in this model, we observed markedly decreased levels of the microglial receptor TREM2, which is implicated in microglial functions including phagocytosis, apoptosis and inflammatory responses following HIV infection and METH treatment. Analysis of our model showed that decreased TREM2 function may lead to HIV- and METH-associated pathological changes. Overall, our assembloid model could be a valuable tool for future analyses of HIV/METH/CNS interactions and mechanisms underlying HAND, which could lead to novel therapeutic approaches to decreasing the CNS viral reservoir.

## INTRODUCTION

Human immunodeficiency virus type-1 (HIV) continues to have a significant neurological impact despite advancements in antiretroviral therapy (ART), notably because the central nervous system (CNS) acts as an HIV reservoir^1^. Such HIV persistence contributes to HIV-associated neurocognitive disorders (HAND), which affect cognitive function, motor skills, and behavior in more than 50% of HIV-positive individuals, regardless of ART^2,3^. Previous studies suggest that methamphetamine (METH) use exacerbates adverse neurological effects of HIV infection by increasing CNS inflammation, disrupting the blood-brain barrier, and enhancing viral replication within the brain ^4–7^. METH is a highly addictive psychostimulant drug that exerts neurotoxic effects via oxidative stress, inflammation, mitochondrial dysfunction, and excitotoxicity, leading to neuronal loss and impaired synaptic plasticity^8–10^. Within the brain, CNS resident immune cells known as microglia can become productively infected by HIV and could be the primary cellular compartment acting as a viral reservoir. Microglia play a key role in neuroinflammation and are implicated in the pathogenesis of HAND and METH-induced neurotoxicity^5,7,11^. However, the lack of human brain-representative models hinders efforts to define relevant mechanisms associated with HIV infection and METH-associated neurotoxicity. Although potential mechanisms underlying HAND have been reported based on analysis of post-mortem brain tissue from subjects with HIV-1-associated dementia (HAD)^12–14^, these approaches cannot address the effects of early infection on disease progression from mild neurocognitive disorder or asymptomatic neurocognitive impairment due to combining HIV infection with METH. Nonetheless, data derived from post-mortem analysis of patients has been validated in simian immunodeficiency virus (SIV)-infected non-human primates, humanized rodent models, and *in vitro* two-dimensional (2D) cellular models^15–20^. However, these methods fail to capture distinct and dynamic aspects of human brain physiology and inter-individual variations. There are currently no effective therapies that address HAND symptoms; thus, creating a suitable experimental model using pertinent human neuronal and microglial cell lineages is critical.

Previous studies have focused on the differentiation of human induced pluripotent stem cells (hiPSCs) into different brain region-specific organoids to serve as models to investigate neurodevelopment, host-pathogen interactions, and neurological disorders^21–27^. However, these widely used brain organoid systems lack microglial cells, which function in neuroinflammatory events in diverse contexts. Therefore, these models cannot accurately recapitulate the neuropathogenesis of METH use in the presence of HIV infection.

Microglia constitute 0.5-17% of total cells in the brain depending on region and play a central role in neuroinflammation^28–30^. Although little is known of their precise role in HIV neuropathogenesis, microglial activation is implicated in the pathogenesis of HAND and METH-induced neurotoxicity^31,32^. Interestingly, TREM2 (Triggering Receptor Expressed on Myeloid Cells 2), a protein expressed on the surface of microglia, regulates activities such as phagocytosis, inflammatory responses, and cytokine/chemokine production in the brain^32,33^. TREM2 also interacts with other proteins and signaling pathways to modulate microglial activity in response to changes in the CNS environment^34^. Some studies suggest that dysregulated TREM2 expression or function in microglia contributes to neuroinflammation, including underlying neurodegenerative disorders^35–37^. However, how TREM2 functions in HAND induction and associated neuroinflammation or neurodegeneration in the presence of METH use is yet to be explored.

Here, we created a next-generation three-dimensional (3D) human brain assembloid model consisting of cerebral organoids (CORGs) and microglia derived from human ESCs/iPSCs. To do so, we developed microglia-like cells using hiPSCs that expressed physiologically relevant markers expressed by *in vivo* microglia. We then used these cells to create a model to support HIV infection and recapitulate hallmarks of microglial activation and proinflammatory cytokine release seen in patients positive for HIV infection and METH use. Specifically, we developed a next-generation 3D assembloid model system consisting of iPSCs/ESCs-derived microglia (iMGs) integrated into well-characterized CORGs. Following HIV infection and METH exposure, cells in the assembloid model showed inflammatory responses, cytokine release, damage to neurons and astrocytes, and impaired synaptic markers—the major hallmarks in the CNS of HIV-infected and METH-exposed patients. Overall, this model could advance understanding of the pathophysiology of HAND and substance use disorders, provide insight into mechanisms underlying neuroinflammation, and lead to targeted interventions to improve brain health and quality of life in vulnerable populations.

## RESULTS

### Generation of induced microglial-like cells from human embryonic stem cell/human induced pluripotent stem cell lines

Model systems exist to study HIV infection of microglia, but their use is limited due to the restricted availability of human microglia from fetal tissue or postmortem brain tissues. Thus, such studies are more often conducted in humanized mouse models or microglial lines such as HMO6 ^38^, HMC3 ^38^, and hTERT-immortalized glial (hT-iMGs) cells ^38,39^, which differ substantively from primary human microglia in morphology, genotype, function, and genetics ^40,41^. Recently, others have differentiated human microglia from iPSC and hESC ^42–44^. Here, to analyze microglia in the context of HIV-associated neurological injury we differentiated human microglia from three different hESC/hiPSC lines using the methodology of McQuade et al. ^45^ (**Figure 1a-c and Figure S1b-c**). Briefly, we generated CD43+ hematopoietic progenitor cells (HPCs) from hESC/iPSC and differentiated them into microglial-like cells (iMGs) in serum-free conditions and in the presence of M-CSF, IL-34 and TGFβ-1 growth factors (**Figure 1a-b**). Flow cytometric data revealed that this protocol yielded iMG cells with high purity (>98.2%,). We then characterized iMGs after 25 days of differentiation by immunocytochemistry for microglial-specific markers, namely Iba-1, P2RY12 and TMEM119 (**Figure 1c and Figure S1b-c**). qRT-PCR showed that iMG cells also express microglial signature genes, namely, Iba-1, CD68, C1QA, PU.1, GPR34, PROS1, CX3CR1 and TMEM119 (**Figure 1d**). As a functional validation, we also found that after lipopolysaccharide (LPS) stimulation, iMGs express genes encoding for the cytokines IL-6 and IL-1β, as anticipated (**Figure 1e**).

**Figure 1:**
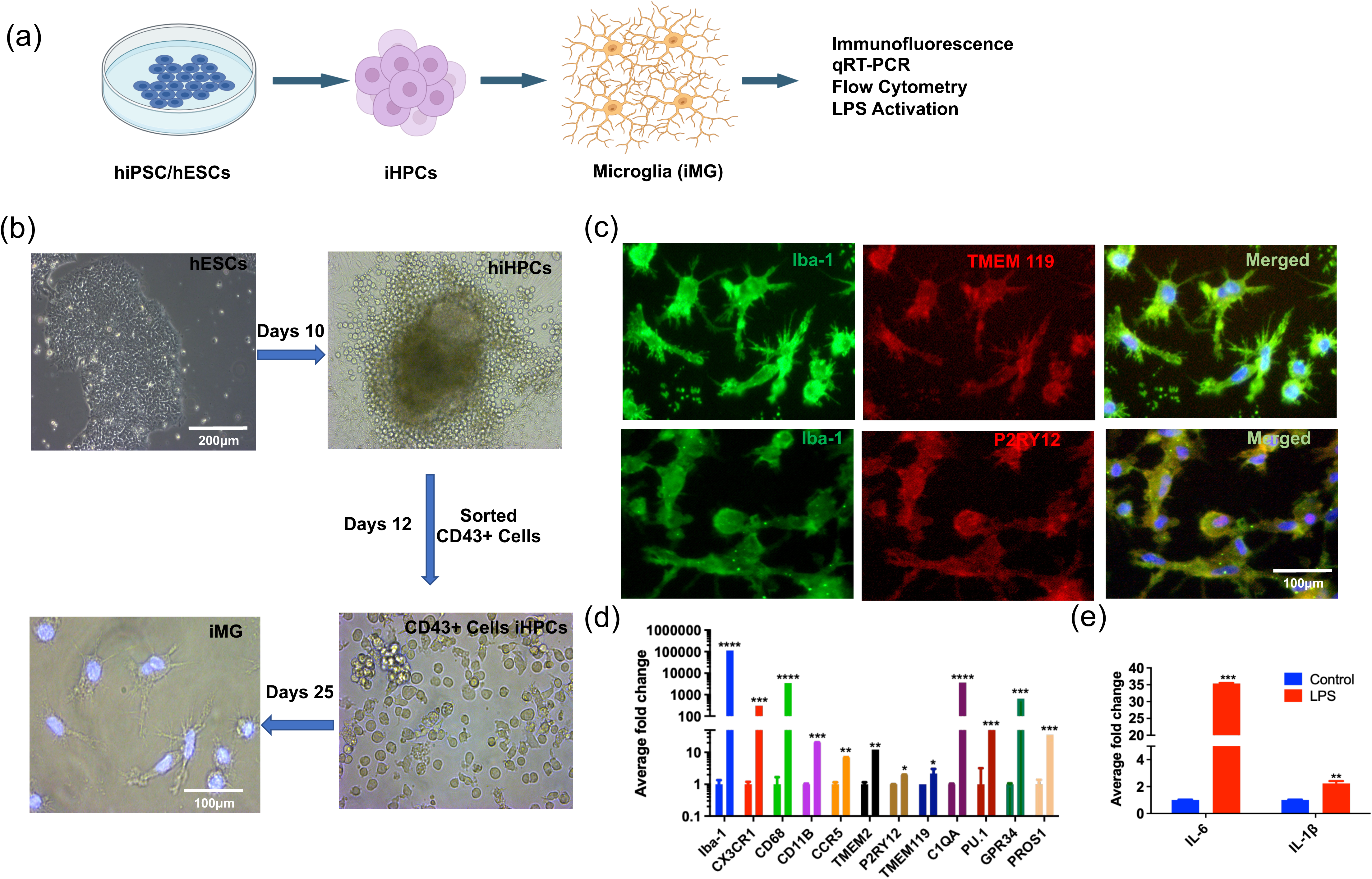
Characterization of human microglial (iMG) cells derived from hESCs/hiPSCs. (a) Schematic showing steps used to generate and differentiate human microglial (iMG) cells from hESCs/hiPSCs. (b) Representative phase contrast images of hESCs differentiated into CD43+ hematopoietic progenitor cells (HPCs) over 10 days and then sorted for CD43+ cells. Cells were cultured in serum-free microglial differentiation medium containing human recombinant proteins M-CSF, IL-34, and TGFβ-1(as described in methods). Scale bar, 100μm. (c) Immunocytochemical analysis showing co-staining of differentiated human microglia with the indicated microglial markers. Merged panels also include the nuclear stain DAPI. Scale bar, 100μm. (d) qRT-PCR analysis of indicated microglial signature genes in iMG cells derived from hESCs. Average fold changes in hESC vs iMG of each gene are shown with the same color bar. ****p<0.0001, ***p<0.0001, **p<0.005, *p<0.01, n=3. Error bars indicate mean±SEM. (e) qRT-PCR analysis of indicated cytokine genes expressed in iMG cells, before and after 24 hr of LPS (100ng/ml) stimulation. ***p<0.0001, **p<0.005, n=3. Error bars indicate mean±SEM.

### Infection and activation of iMG cells by HIV-1

We next assessed the activity of hESC/iPSC-derived iMG cells after infection with HIV-1. Specifically, we utilized the HIV-1_BaL_ strain, an R5-tropic virus that is typically more efficient at infecting macrophages compared to other strains. To do so, we infected iMG cells with HIV-1_BaL_ (labeled as HIV-1) and then measured the rate of HIV replication over several days by ELISA for the viral protein p24 (see methods). We observed that iMG cells supported HIV-1 replication time-dependently after infection (**Figure 2b,c**) We also infected three different iMG lines with HIV-1 and performed immunocytochemistry to assess potential colocalization of p24 (red) with the microglial marker Iba-1 (green) (**Figure 2c and Figure S2a**). At 12 days post-infection, we observed that >90% of iMG (green) cells were also p24-positive (**Figure 2c and Figure S2a**). We also analyzed iMG infection by HIV-1 at 2 different MOIs at day 12 by evaluating p24 protein levels, indicating that an MOI of 0.5 was sufficient for successful viral infection that further increased at MOI of 2 (**Figure 2d**). Moreover, HIV infection also led to an activated phenotype in microglia, as indicated by IL-6 release into the supernatant up to day 9, followed by a decrease at day 12 (Figure 2e). We next used purified HIV proteins gp120 and Tat and treated iMG cells with each protein individually. By 12 hours of treatment, we observed microglial activation after treatment with either Tat or gp120 protein, based on IL-6 release (**Figure S2c**). Next, we treated neurons differentiated from human neural stem cells (hNSCs) with conditioned medium from HIV-infected iMG cells that may contain HIV virus, HIV proteins and/or secreted cytokines. We found that conditioned medium from HIV-infected iMG cells enhanced neuronal apoptosis and reduced the length and size of neuronal fibers compared to mock or control groups (**Figure S2d**). Overall, we conclude that iMG cells infected with HIV-1 become activated and release cytokines that may injure neurons.

**Figure 2:**
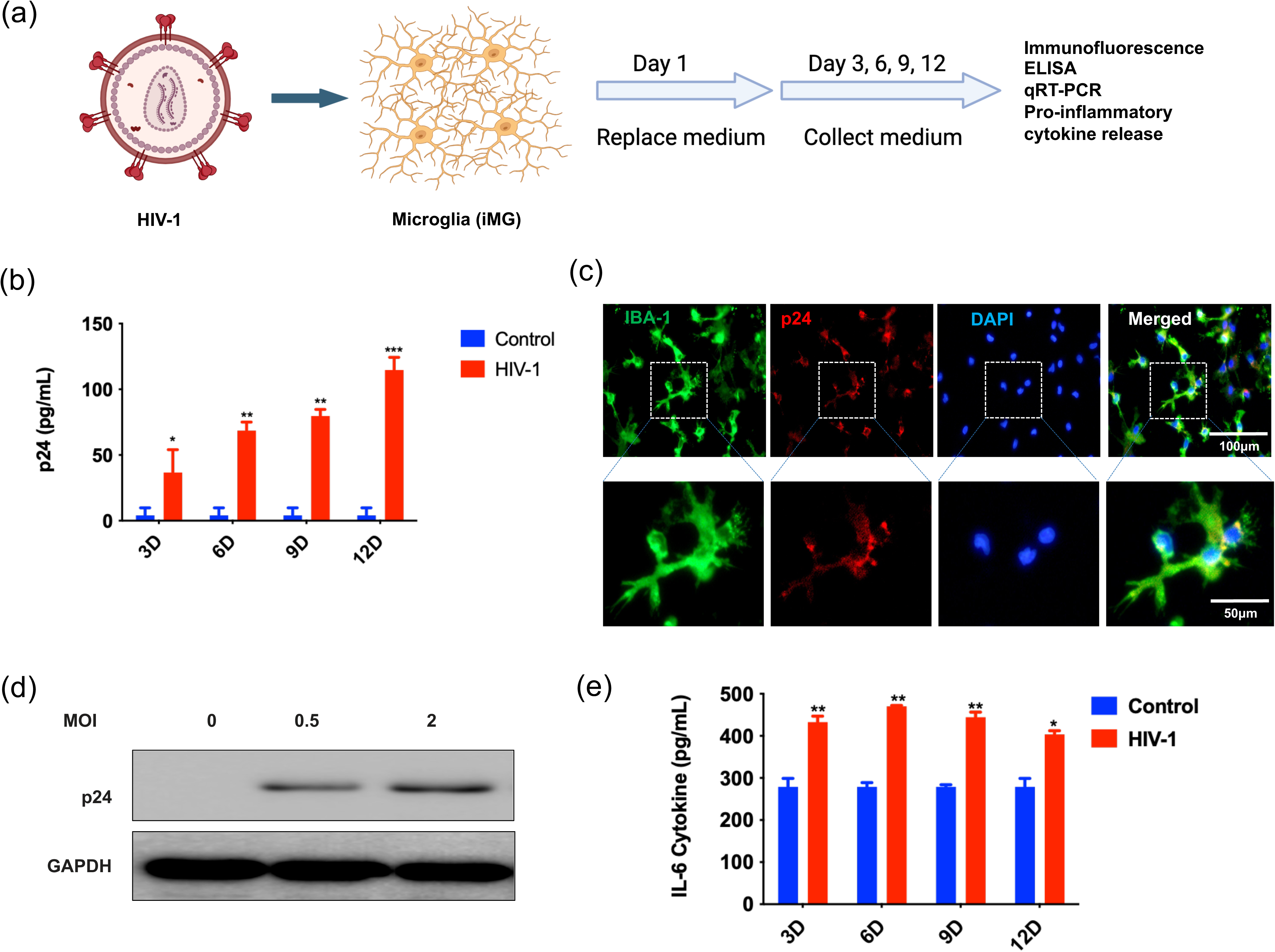
HIV-1 infection of iMG cells activates microglia and promotes cytokine release. (a) Schematic showing HIV-1 infection of iMG cells at indicated time points and characterization of infected cells as shown in panels below. (b) iMG cells support HIV-1replication, analyzed by p24 ELISA of culture supernatants. **p<0.005, *p<0.01, n=3. Error bars indicate mean±SEM. (c) (Top row) Immunocytochemical analysis of iMG cells infected with HIV-1 and co-stained for the microglial marker Iba-1 (green), the HIV protein p24 (red), and the nuclear stain DAPI. (Bottom row) Higher magnification images of corresponding boxed areas in the panel above. Scale bar: 100μm (d) Representative immunoblot for the viral protein p24 of iMG cells infected with HIV-1 at indicated MOIs. GAPDH served as a loading control. n=3. (e) Secretion of the cytokine IL-6, as detected by ELISA analysis of supernatants from control and HIV-1_BaL_-infected iMG cells. Cells were infected on day 0 and assessed at indicated time points. Control vs HIV-1_BaL_, *p<0.01, **p<0.005

### The 3D cerebral assembloid model (3D-CBAM) exhibits diverse CNS cell types that participate in crosstalk as seen in HIV-1 and METH neuropathogenesis

Post-mortem analysis of brain tissues from individuals who were both METH users and infected with HIV-1 shows neuroinflammation and viral infection of human macrophages and microglia^29,46–49^. Cerebral organoids have been used to analyze comparable CNS conditions, as organoids contain diverse cell types, including neural stem cells, oligodendrocytes, neurons, and astrocytes. However, these organoids lack microglia, which are relevant to conditions such as HAND. Thus, to create an organoid system that includes microglia we co-cultured CD43+ HPCs for 15 days with cerebral organoids derived from three different hESC/iPSC lines, as described with slight modifications ^42^. At the end of the culture period, CD43+ cells were distributed throughout and incorporated within organoids and extended their ramified processes, characteristic of microglia to varying extents within the 3D cerebral organoid environment (**Figure 3a-b and Figure S3a-b**). To characterize the maturation of cells in the assembloids, 3D-CBAMs from three stem cell lines were harvested at 120 days, sectioned, and stained for neuronal and glial markers. Immunohistochemistry analysis revealed cells positive for the neuron-specific cytoskeletal marker β-III-tubulin and the neuro-progenitor marker SOX2, in addition to GFAP+ and Iba-1+ glial cells (**Figure 3c and Figure S3 a-b**). Further, fluorescence microscopy analysis of tissue sections revealed dense neural and glial networks within 3D-CBAMs (**Figure 3b,c and Figure S3 a-b**). We also found that 3D-CBAMs, relative to CORGs which lack microglia, significantly expressed microglial signature genes such Iba-1, CX3CR1, CD68, CD11b, C1QA, PU.1, PROS1, GPR34, P2RY12, and TMEM119 (**Figure 3d**).

**Figure 3:**
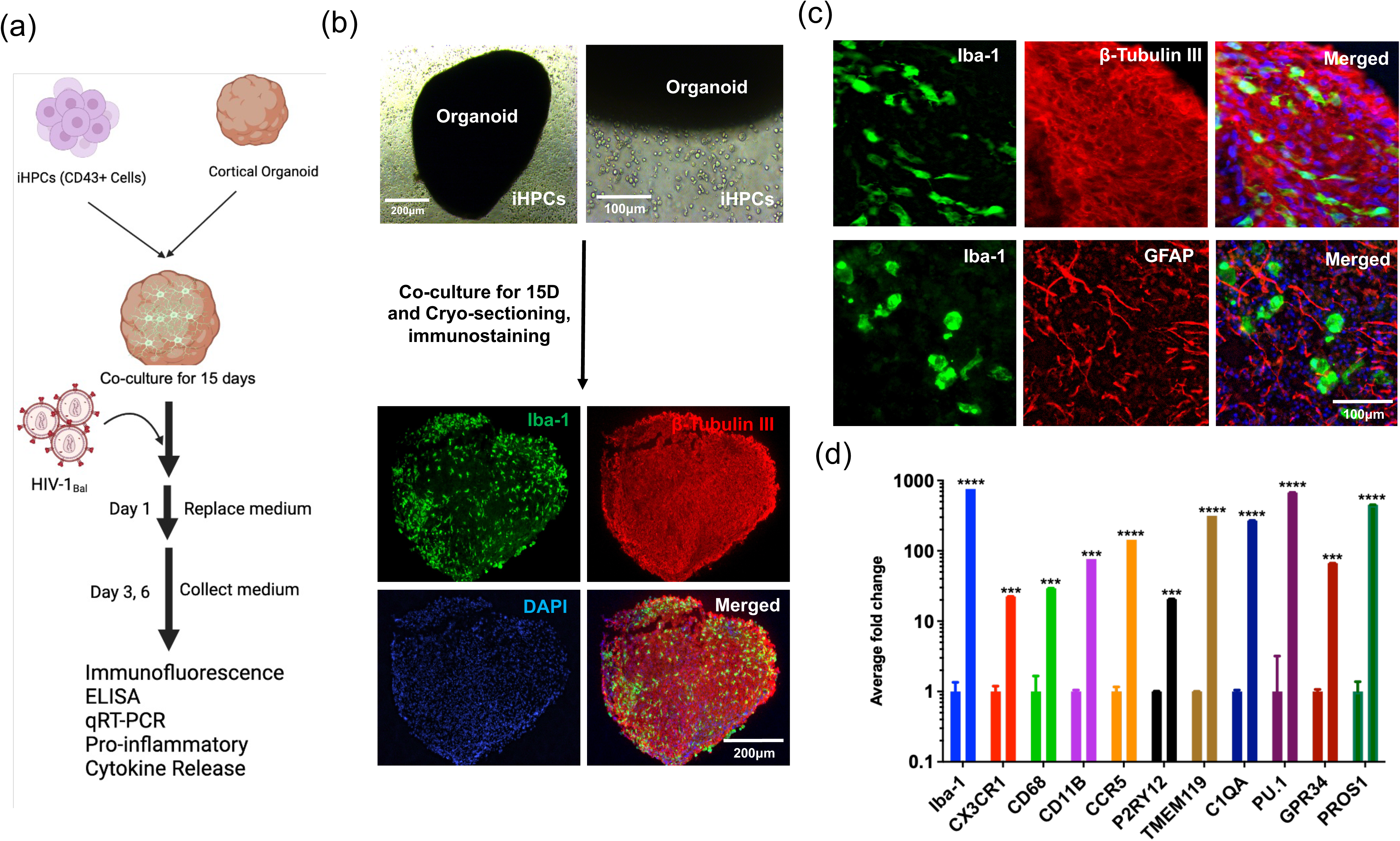
Generation of the 3D cortical assembloid model. (a) Schematic showing generation of the 3D cortical assembloid model in which iMG cells are integrated into cerebral organoids. (b) (Upper) Representative phase contrast images of co-cultures of CD43+ cells and cerebral organoids. CD43+ cells become attached to the organoid. (Lower) Immunohistochemical analysis of co-cultures after 15 days with indicated microglial (Iba-1, green) and neuronal (beta-tubulin-III, red) markers, plus the nuclear stain DAPI. Image indicates that hiMGs are evenly integrated throughout the organoid and extend ramified processes. Scale Bar: 200μm. (c) Representative higher magnification images of co-cultures described in (b, lower) showing labelling with microglial (Iba-1, green), neuronal (beta-tubulin-III, red) and astrocytic (GFAP, red) markers. Scale Bar: 100μm. (d) qRT-PCR analysis showing the comparison of microglial markers expression in 3D organoid, and 3D cortical assembloid model. Cortical organoid (CORG) vs assembloid (CBAM) compares the average fold change with respective of each gene of same color bar graph ****p<0.0001, ***p<0.0001, **p<0.005, *p<0.01, n=3 Error bars indicate mean± SEM

HIV and METH-associated neuropathogenesis generally impact glutamatergic neurons^50^. Thus, we used immunofluorescence analysis to search for glutamatergic neurons in 3D-CBAM cultures and found that most differentiated neurons expressed VGLUT1+ (vesicular glutamate transporter 1) (**Figure S4a**). We also evaluated for expression of the pre-synaptic protein synaptophysin (SYN) and the post-synaptic marker PSD95 in 3D-CBAM cultures (**Figure S4b-c**). In these cultures, we also observed puncta of colocalized SYN and PSD, indicating synaptic connectivity (**Figure S4c**). Collectively, our results indicate that our 3D-CBAM model is a representative *in vitro* brain model system potentially useful to assess the effects of HIV-1 infection in the presence of METH.

### HIV-1 infection of a cerebral assembloid model containing iMG cells promotes cytokine release

We next asked whether HIV-1_BaL_ infection of microglia in a 3D-CBAM model recapitulates features associated with HIV neuropathogenesis in human brain. To do so, we established a 3D-CBAM model containing HIV-1_BaL_-infected iMG infection as described above, performing immuno-labelling for both a microglial (Iba-1) and viral (p24) marker (**Figure 4a**) which showed co-labeling of iMGs with HIV-1 protein p24. Interestingly, we also observed an increased release of viral particles into culture supernatants over several days (**Figure 4b**), indicating that the model supports HIV-1_BaL_ infection and replication. Viral infection of 3D-CBAM was confirmed by immunoblotting for p24 (**Figure 4c**). We conclude that our model system supports HIV-1_BaL_ infection by specifically targeting the microglial population.

**Figure 4:**
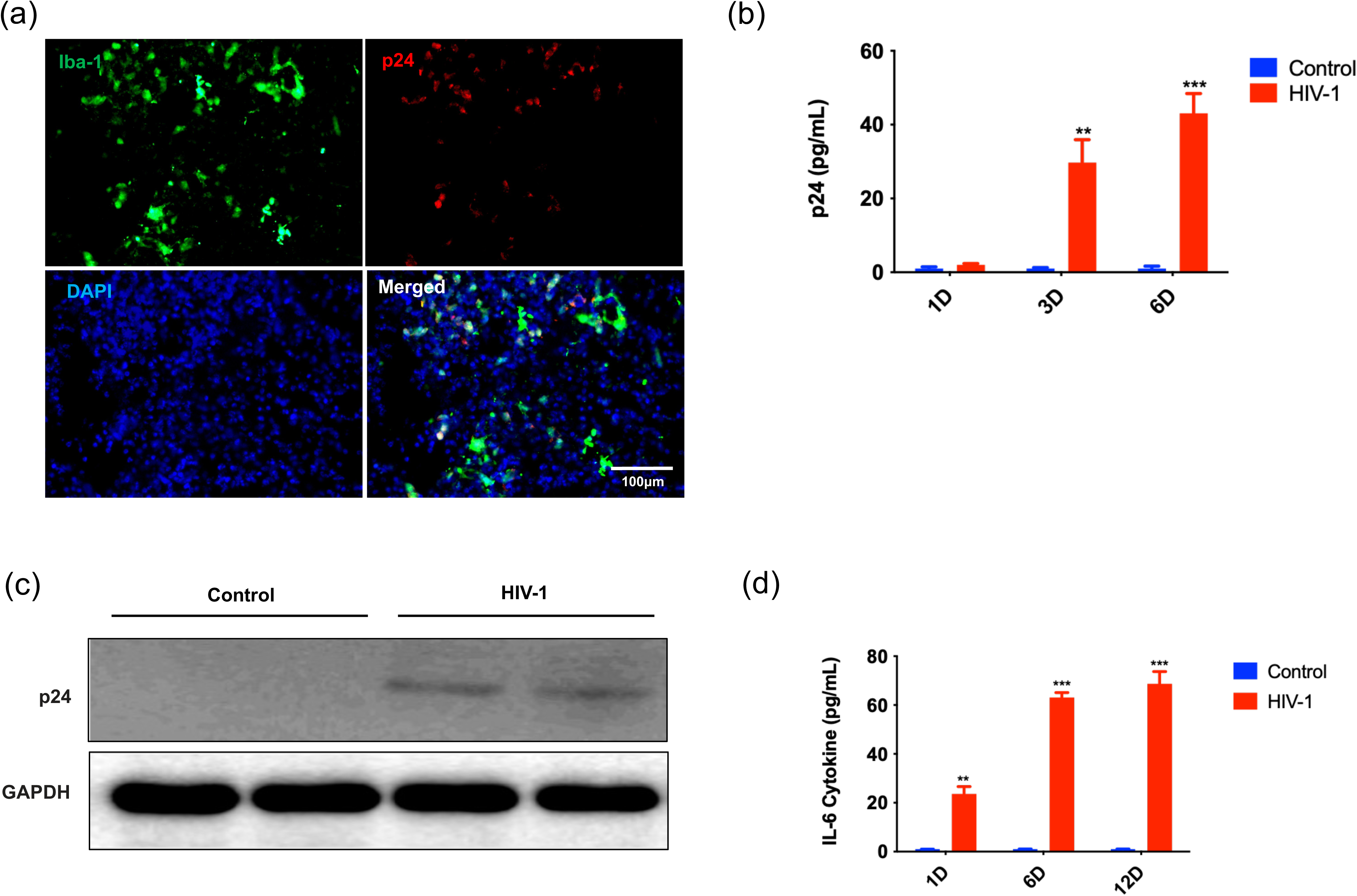
HIV can infect the 3D cerebral assembloid model, propagate, and promote IL-6 release. (a) Immunocytochemical analysis with the indicated markers of 3D cerebral assembloids following HIV-1 infection of iMG cells. Shown are representative images. Scale bar: 100μm. (b) Culture supernatants were collected from HIV-1_BaL_-infected 3D assembloids at indicated days after infection and analyzed by ELISA for the viral protein p24. Results indicate replication of HIV-1. Control *vs* HIV **p<0.001. n=3, mean± SEM. (c) Representative immunoblot for p24 in HIV-1_BaL_-infected 3D assembloids. GAPDH served as a loading control. n=3 (d) Cytokine IL-6 secretion by iMG-containing 3D cerebral assembloids at indicated time points after infection with HIV-1virus, based on human IL-6 ELISA analysis of culture supernatants. Control vs HIV-1**p<0.001.

Activated microglia and macrophages associated with HIV neuropathogenesis reportedly release the pro-inflammatory cytokines IL-6 and IL-1β, which function in neuroinflammation^51,52^. Thus, we performed ELISA to measure IL-6 levels in supernatants from 3D-CBAM. Supernatants from HIV-1_BaL-_infected 3D-CBAM showed significantly higher levels of IL-6 than the mock-infected control group, and those levels increased until day 12(**Figure 4d**).

Overall, these results strongly suggest that our 3D-CBAM model supports chronic HIV replication and recapitulates a neuroinflammatory phenotype comparable to that observed in HIV-infected patient brain samples.

### METH treatment potentiates HIV-1 infection and replication in iMG cells and in a 3D cerebral assembloid and activates cytokine release by glial cells

HIV infection and METH use synergize to promote neurotoxicity, potentiate neuroinflammation, disrupt neurotransmitter activity, and compromise neuroplasticity, contributing to HAND^2,48^. Thus, we asked if the presence of METH enhances HIV infection in our iMG (Figure 5a-c) and 3D cerebral assembloid (Figure 5d-e) by infecting hiPSC-derived iMGs with HIV-1_BaL_ in the presence and absence of METH. Interestingly, METH treatment increased HIV-1_BaL_ infection of iMG cells as compared to cultures not treated with METH, as indicated by co-labelling of HIV marker p24 and the microglial marker Iba-1. Moreover, p24 levels increased in culture media over several days of culture, suggesting that METH enhances viral replication (**Figure 5a-b**). Similarly, METH treatment plus HIV infection led to an activated phenotype in microglia, indicated by IL-6 release into the supernatant until day 12 (**Figure 5c**). Next, we infected 3D-CBAMs with HIV-1_BaL_ for 12 days and collected supernatants for ELISA analysis to assess cytokine release and virus replication. Assembloids containing microglia supported HIV replication over time, and METH treatment enhanced that replication, as quantified by p24 levels (**Figure 5d**). HIV infection plus METH treatment also enhanced the release of the pro-inflammatory cytokines IL-6, TNF-a, and IL-1β into the supernatant compared to non-treated NT control (**Figure 5e**). We next asked if METH and HIV infection affect the glial cell population in the assembloid model by immunocytochemical analysis of sections of assembloids with microglial (Iba-1) and astrocytic (GFAP) markers. As expected, METH treatment plus HIV infection led to an ameboid morphology in microglia and resulted in activation of microglia and astrocytes, which may release cytokines and exacerbate neurodegeneration (**Figure 5f**) ^53,54^. Overall, these findings confirm that 3D-CBAM is an appropriate model system for examining mechanisms underlying neurological disorders associated with HIV infection and drug abuse.

**Figure 5:**
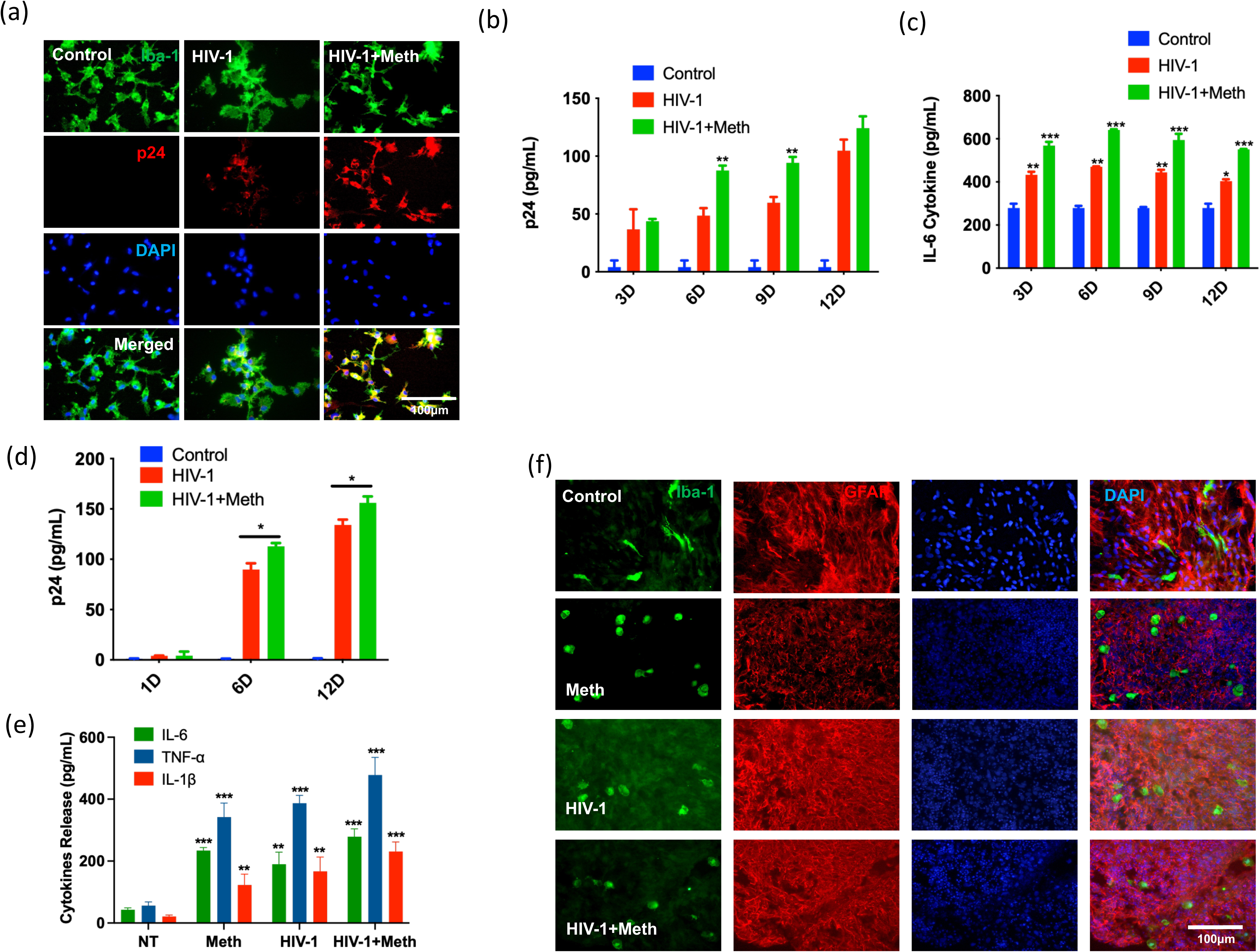
METH treatment of 3D cerebral assembloids containing iMG cells potentiates HIV-1 infection and replication and activates cytokine release. (a) Immunocytochemical analysis of control, HIV-1_BaL_-infected, and HIV-1_BaL_-infected/METH-treated iMG cells for Iba-1 (microglial marker, green) and p24 (HIV marker, red). Scale bar: 100μm (b) Quantitative analysis of p24 ELISA over a 12-day time-course after infection of experimental groups shown in (a). Data suggests that iMG cells support HIV-1 replication and that METH treatment enhances HIV-1 replication. **p<0.005, *p<0.01, n=3. Error bars indicate mean± SEM (c) Secretion of cytokine IL-6 by cells described in (b). IL-6 was assessed by ELISA of culture supernatants. Control vs HIV-1 *p<0.01 (d) Quantitative analysis of p24 ELISA over a 12-day period after HIV-1 infection of 3D cortical assembloids, with or without METH treatment. Shown is analysis of supernatants from control, HIV-1_BaL_-infected, or HIV-1_BaL_-infected/METH-treated assembloids at indicated days after treatment. **p<0.005, *p<0.01. n=3. Error bar indicates mean±SEM. (e) Secretion of indicated cytokines by untreated (NT) control, METH-treated, HIV-1-infected, or HIV-1-infected/METH-treated assembloids. Cytokines were assessed by ELISA of assembloid supernatants. Control vs HIV-1*p<0.01 (f) Immunofluorescence analysis of control, METH-treated, HIV-1-infected, or HIV-1-infected/METH-treated assembloids stained for microglial (Iba-1+) and astrocytic (GFAP+) markers plus DAPI. Scale bar: 100μm.

### HIV-1 infection and METH treatment of 3D cerebral assembloids containing iMG cells decreases TREM2 expression and enhances specific lincRNAs

Neurotoxic mechanisms associated with HAND and METH-associated neuropathogenesis promote microglial activation^55–57^. The receptor protein TREM2 is predominantly expressed on the surface of activated microglia, where it regulates functions associated with phagocytosis and inflammatory responses. To determine whether TREM plays a role in HIV/METH comorbidity associated HAND we explored the effects of HIV infection on the activity of TREM2 and the complement system in iMG cells. Interestingly, we observed decreased expression of TREM2 and DAP12 receptors relative to expression of the complement genes C3a and C1q in microglial cells after HIV-1 infection (**Figure 6a).** Reduced TREM2 levels were confirmed by immunoblotting and co-labeling of TREM2 with the microglial marker P2RY12 in the HIV-infected microglial group (**Figure 6b-d**. Next, we assessed TREM2 in the HIV-infected assembloid model compared to an uninfected control group and observed significant TREM downregulation (**Figure 6e**). Interestingly, TREM2 levels were lower only in cerebral organoids lacking microglial cells; however, assembloids with microglia showed higher TREM2 expression levels, confirming that TREM2 is expressed by microglial cells (**Figure 6e**). These results indicate that combined HIV/METH treatment activates microglia. Thus, we examined HIV and METH effects on TREM2 levels. mRNA and protein analysis confirmed that HIV/METH co-treatment significantly decreased TREM2 expression relative to the control group (**Figure 6f-h**), suggesting that dysregulated TREM2 expression or function in microglia may contribute to neuroinflammation seen in HIV infection and METH neurotoxicity.

**Figure 6:**
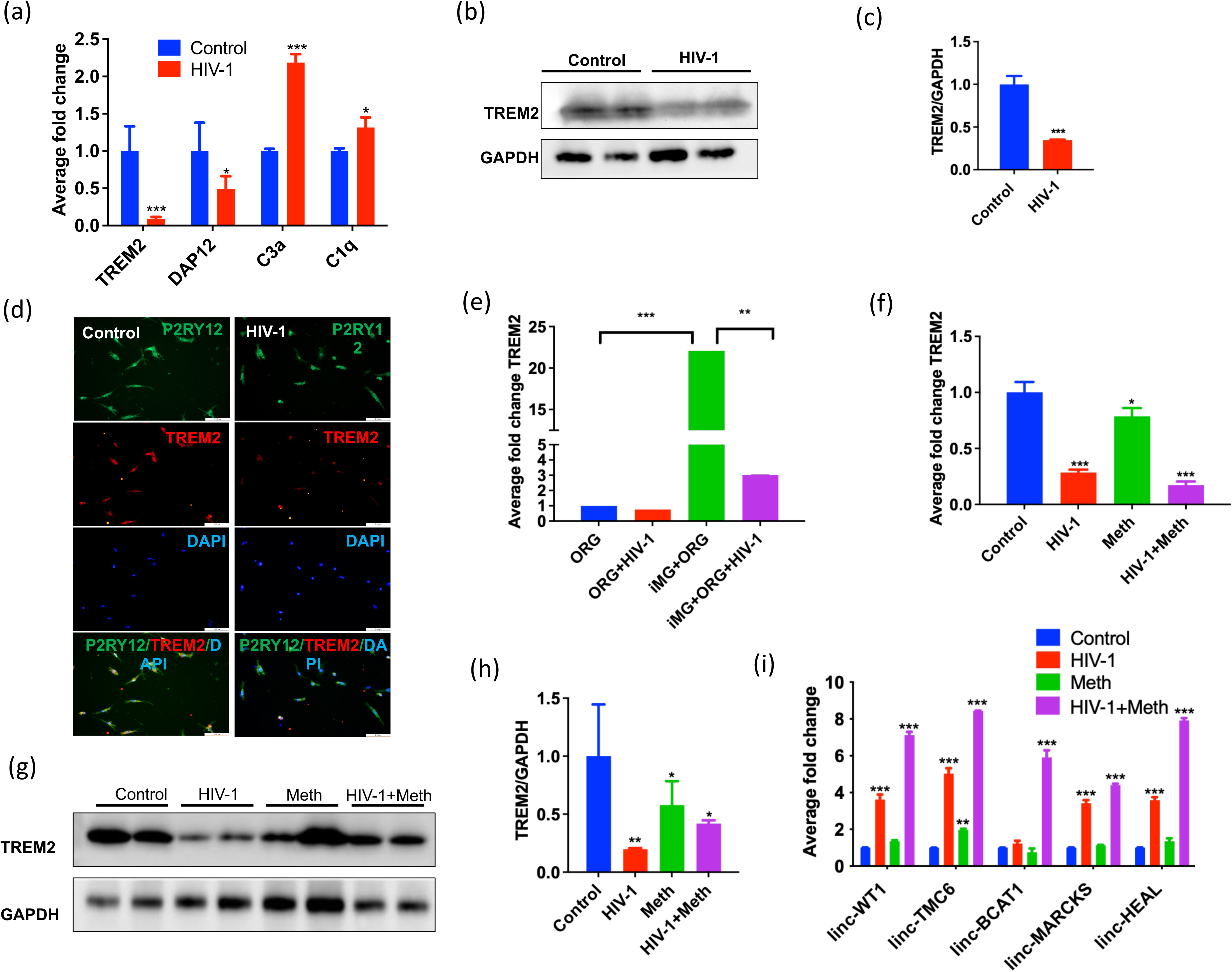
TREM2 expression decreases in HIV-1-infected, METH-treated iMG cells and 3D cerebral assembloids. (a) qRT-PCR analysis showing expression of TREM2 as well as complement system genes in control and HIV-1_BaL_-infected iMG cells. ***p<0.0001, *p<0.01. n=3, Error bars indicate mean± SEM. (b) Representative immunoblot for TREM2 protein in control and HIV-1-infected iMG cells. (c) Bar graph showing quantification of TREM2 protein level normalized to GAPDH in control and HIV-1 infected iMG cells, ***p<0.0001, n=3. Error bars indicate mean± SEM. (d) Immunofluorescence images showing control and HIV-1_BaL_-infected iMG cells stained for TREM2 (red), the microglial marker P2RY12 (green), and DAPI. n=3. Scale bar: 100μm. (e) qRT-PCR analysis of TREM expression in organoids only (ORG), and organoids infected with HIV-1(ORG-HIV-1), treated with METH (METH) or both (HIV-1_+_ METH). ***p<0.0001, **p<0.001. n=3. Error bars indicate mean± SEM. (f) qRT-PCR analysis showing the TREM2 expression in the 3D cortical assembloid model treated as described in (d). ***p<0.0001, **p<0.01. n=3. Error bars indicate mean± SEM. (g) Representative immunoblot of TREM2 protein in indicated groups of the 3D cortical assembloid model. (h) Bar graph showing quantification of TREM2 protein level normalized to GAPDH in control, HIV-1, Meth and Meth+HIV-1 in assembloid model, **p<0.001, *p<0.01, n=3. Error bars indicate mean± SEM. (i) qRT-PCR analysis showing expression of indicated lincRNAs associated with HIV infection in assembloid models of indicated groups. ***p<0.0001, **p<0.001. n=3. Error bars indicate mean±SEM.

We previously reported that TLR stimulation and HIV infection regulate lincRNAs such as lincWT1A, TMC, BCAT, MARC and HEAL in HIV-infected macrophages and PBMCs ^58–60^. Thus, we asked whether HIV infection and/or METH exposure regulate lincRNA expression in iMG cells. In most cases, exposure to METH alone had little effect on lincRNA expression, while HIV infection upregulated lincRNA expression, and combining METH and HIV infection significantly upregulated expression of all lincRNAs tested at day 12 (**Figure 6i**), confirming that HIV infection regulates expression of several lincRNAs in microglia.

### HIV infection in the 3D-CBAM model mimics CNS pathology in HIV-infected post-mortem brain tissue reports

Finally, to determine whether HIV infection or METH treatment and resultant inflammatory conditions in infected 3D-CBAMs would alter the activity of neurons, as the cell type most susceptible to damage due to HIV, we evaluated neuronal cell death in sections of 3D-CBAMs by immunostaining with markers of apoptosis (cleaved-caspase3) and neurons (MAP2) (**Figure 7a-b**). We observed an increase in the percentage of cells labeled by both cleaved-caspase and MAP2 cells in 3D-CBAMs exposed to HIV-1 infection and METH treatment relative to control 3D-CBAMs (**Figure 7b**). Moreover, neuronal fibers were of normal size and length and showed healthy density in the control group but were reduced in METH, HIV, and HIV+METH treated groups, as shown by co-labeling with neuronal marker MAP-2 (**Figure 7a**). These results indicate that HIV-1 infection may enhance neuronal loss in the 3D-CBAM model. Next, we assessed potential synaptic damage following HIV infection or METH treatment in the 3D-CBAM model by labeling cells in organoid sections with the pre-synaptic marker synaptophysin (SYN) (**Figure 7c**). We observed a marked reduction in SYN levels in METH, HIV, and HIV+METH treatment conditions relative to untreated controls, suggesting that pre-synaptic terminals are compromised by HIV-1 infection and METH treatment relative to controls. Overall, these findings support the physiological relevance of the 3D-CBAM system to the analysis of HIV-1 and METH neuropathogenesis *in vitro*.

**Figure 7:**
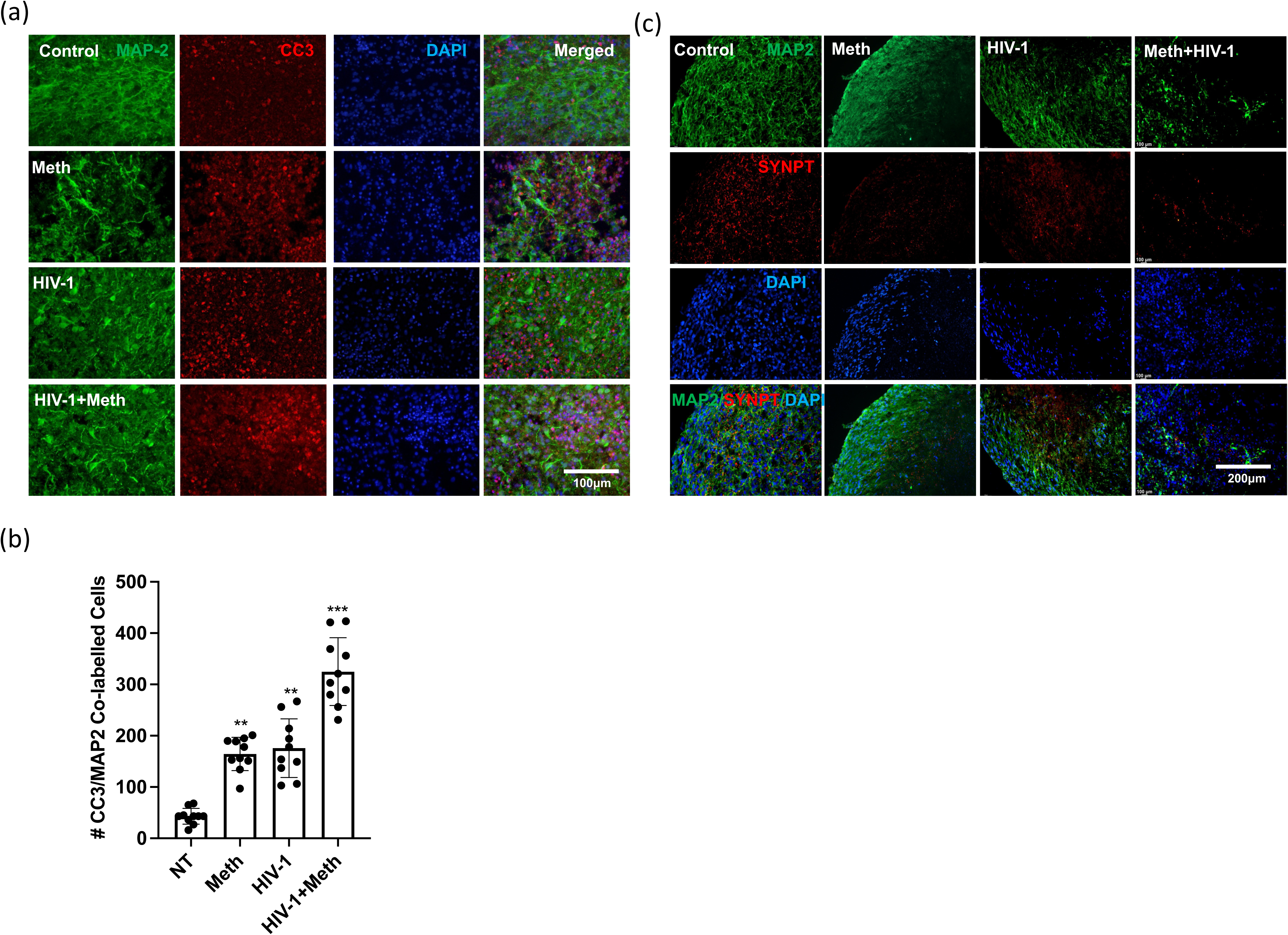
HIV infection of the 3D cerebral assembloid model promotes decreases in levels of neuronal and synaptic markers. (a) Immunohistochemical analysis of tissue sections from the cerebral assembloid model in control, HIV-infected, METH-treated, or HIV-infected/METH-treated groups stained for the neuronal marker MAP-2 (green), the apoptotic marker cleaved caspase-3 (red), and DAPI. n=3. Scale bar: 100μm. (b) Quantitative analysis of samples shown in (a). Shown is the ratio of cleaved caspase-3+ staining to MAP-2 staining in indicated treatment groups in ten different section’s view. n=3. ***p=0.0001, **p<0.001. n=3. Error bars indicate mean± SEM. (c) Immunohistochemical analysis of sample groups shown in (a) stained for MAP-2 (green), synaptophysin (SYNPT, red), and DAPI. Data indicates that METH treatment and HIV-1infection may decrease levels of SYN protein and size of neuronal fibers. Scale bar 100μm.

## Discussion

To investigate HIV- and METH-associated neuropathogenesis, we developed a brain region-specific 3D *in vitro* cerebral assembloid model (3D-CBAM) derived from human pluripotent stem cells that recapitulates the neuroinflammatory microenvironment seen in human brain pathophysiology. To do so, we differentiated iMG cells from hESCs/hiPSCs using an established protocol with minor modifications ^45^ and also generated and characterized CORGs containing neural progenitor cells, astrocytes, and neurons using our previously established method based on a Matrigel-embedded system. We then generated a 3D cortical assembloid system by integrating iMGs into CORGs and appiled this system as a model to investigate HIV- and METH-associated pathologies.

Analysis of HAND-associated neurological responses and neurodegeneration *in vitro* is challenging due to the complexity of the CNS and its diversity in cellular interactions. Other studies report establishing neurons, astrocytes, brain organoids, and microglia using hiPSCs, shedding light on our understanding of host-HIV interactions^25,61–63^. Recently various immortalized microglial lines have been developed that can be expanded indefinitely but differ from cells found in *in vivo* conditions^64^. Primary microglia from embryonic and adult brain tissue show a limited life span in culture. Overall, studies of HIV infection and METH exposure are hindered by limited tissue availability and variability of patient samples. Previous studies have incorporated microglia into organoids to investigate neurodegenerative and inflammatory disorders ^25,61–63^, but mechanisms underlying HIV infection and drug use are elusive. Here, for the first time, we report the creation of a 3D cortical assembloid model (3D-CBAM) to evaluate the comorbidity of HIV infection and METH exposure and its effect on mechanisms underlying neuroinflammation and neurotoxicity. Our findings show that incorporating iMG cells into a 3D-CBAM model that has been infected by HIV replicates conditions associated with HIV infection *in vivo*.

An important feature of HIV- and METH-associated neuropathology is microglial activation and release of pro-inflammatory cytokines^65,66^. In our 3D-CBAM model system, we observed that HIV infection and METH exposure rapidly induce the secretion of cytokines such as IL-6, IL-1β and TNF-α, which play a major role in neurodegeneration and CNS injury. The increase in levels of these cytokines over time was positively correlated with HIV replication, a finding consistent with prior studies of patient brain samples and microglial cell lines ^4,67^. Pro-inflammatory cytokines released by infected microglial cells also contribute to the release of other soluble neurotoxic factors contributing to neurodegeneration and HAND-like symptoms^68,69^. In our model system, we observed morphological changes in astrocytes in the 3D-CBAM model system following METH exposure and HIV infection, as well as enhanced neuronal cell death and a reduction in synaptic proteins, based on analysis of apoptotic and synaptic markers. These outcomes further supported by prior study highlighted METH and HIV enhances the autophagy and apoptotic cell death in astrocytes^50^.

Recently, the TREM2 receptor expressed on microglia has emerged as an important regulator of neuroinflammation, clearance of extracellular proteins, and neuroprotective pathways^70,71^. TREM2 dysregulation is also associated with neurodegenerative disorders^72–74^. However, its role in HIV- and METH-associated neuropathogenesis has not been studied. In our model, we showed that HIV infection and METH exposure downregulate cellular TREM2 receptor expression, enhancing neuroinflammatory responses and neuronal injury. Our results corroborate an earlier finding correlating TREM2 expression in brain tissue of HIV patients with HAND^75,76^. Interestingly, the level of soluble TREM2 in CSF increased with systemic and brain HIV-1 disease severity, with the highest levels found in patients with HAND indicating activation of macrophage/microglial cells ^77,78^. Thus, based on our study we propose that decreased cellular TREM2 expression may enhance the release of inflammatory cytokines that underlie neurotoxicity. However, detailed molecular mechanisms underlying TREM2 activity in the context of HIV and substance use disorders require further investigation.

In summary, we report that an assembloid model incorporating iMG cells recapitulates HIV/METH-associated neuropathology and could provide a powerful *in vitro* tool to investigate the molecular dynamics of METH and HIV neuropathogenesis and its progression. This model also provides an excellent platform to understand host-pathogen interactions, drug abuse-associated neurotoxicities, and molecular mechanisms underlying neuropathogenesis and development of HAND. To the best of our knowledge, ours is the first study using a 3D-CBAM model to assess HIV- and METH-associated neuroinflammation and neurotoxicity. Finally, targeting TREM2 in neuroinflammation could represent a promising therapeutic approach to mitigate neurological consequences of HIV infection and METH abuse.

## MATERIALS AND METHODS

### hESC/iPSC culture and maintenance

All studies were conducted in accordance with approved IRB protocols of the University of California, San Diego. H9 human embryonic stem cells (hESCs, WA09) from WiCell or human induced pluripotent stem cells (hiPSCs, CVB from Coriell also termed GM25430 and EC1 ^79^ were cultured in 6-well plates (Corning) in feeder-free conditions using growth factor reduced Matrigel (MTG, BD Bioscience) in complete MTeSR-1 medium (STEMCELL Technologies) in a humidified incubator (5% CO_2_, 37^0^C), as previously established in our laboratory ^21,22,26,27^. hESCs/hiPSCs were fed fresh media daily and passaged every 7-8 days.

### Cerebral organoid generation

Human 3D cerebral brain organoids were generated using a previously established protocol ^80^. hESCs/hiPSCs were cultured and maintained in 6-well Matrigel-coated tissue culture plates in MTeSR-1 medium in a CO_2_ incubator at 37°C. hESC/hiPSC colonies in cell dissociation medium (STEMCELL Technologies) were gently detached from wells, and colonies were dissociated for 10 min in Accumax solution at 37°C to generate a single-cell suspension. On day 0, embryoid bodies were formed using the hanging drop method with 4500 cells/drop in DMEM/F12 media supplemented with 20% knockout serum replacement, 4ng/ml bFGF, NEAA and glutamine or grown in microwell plates. After 2 days of hanging drop culture, embryoid bodies were transferred to sterile petri dishes with fresh media. After 6 days, embryoid bodies were transferred to new petri dishes containing neural induction media consisting of DMEM/F12, 1:100 N2 supplement, NEAA, glutamine and 1ug/ml heparin until day 11. Then, organoids were transferred to Matrigel droplets and cultured in a 1:1 mixture of DMEM/F12 and Neurobasal medium supplemented with 1:100 B27 without vitamin A, 1:200 N2, NEAA, insulin, β-mercaptoethanol and glutamine. On day 15 organoids were transferred to stir flask bioreactors to allow long-term growth in differentiation media containing retinoic acid and vitamin A. Media was changed every 3 days.

### Differentiation of human induced microglial (iMG) cells

Differentiation of human induced microglial (iMG) cells from hESCs or iPSCs was performed using a previously established 2-step protocol ^45^ with slight modifications) and an HPC differentiation kit (STEMCELL Technologies). In step 1, induced hematopoietic progenitor cells (iHPCs) were generated from hESCs. Briefly, iPSCs were sparsely plated in iPS-Brew with 10 mM ROCK inhibitor (STEMCELL Technologies #72304) onto Cultrex Basement Membrane Matrix-coated 6-well plates using ReLeSR (STEMCELL Technologies #100-0484). Cells were differentiated to CD43+ hematopoietic progenitors using the StemCell Technologies Hematopoietic Kit (STEMCELL Technologies #05310). On day 1, the media was removed, and 1X Supplement A was added. An additional 1 ml/well of 1X Supplement A was added on Day 3, and on Day 5 the cells were changed to 1X Supplement B. Cells received an additional 1 ml/well of 1X Supplement B on Days 7, 9, and 11. The resulting nonadherent hematopoietic progenitor cells were collected on day 12. In step 2, iHPCs were replated onto Cultrex Basement Membrane Matrix-coated plates in microglia media consisting of DMEM/F12 (Gibco #11320033), 2X insulin-transferrin-selenite (Gibco #41400045), 2X B27 supplement (Gibco #17504044), 0.5X N2 supplement (Gibco #17502048), 1X GlutaMAX (Gibco #35050061), 1X Non-Essential Amino Acids (Gibco #11140050), 800 μM monothioglycerol (Sigma-Aldrich #M6145), and 5 mg ml^-1^ insulin (Sigma-Aldrich #I9278) supplemented with M-CSF (25 ng/ml), IL-34 (100 ng/ml; PeproTech), and TGFß-1 (50 ng/ml; Militenyi) made fresh every time. CD43^+^ cells were plated in complete medium at a density 1-2 x10^5^ cells in 2 mL per well in Matrigel-coated 6-well plates. Every 2 days cells were supplemented with 1ml per well of complete iMG differentiation medium. After 25 days of microglial differentiation, we used these cells for HIV infection and treatment with LPS.

### Generation of 3D cortical assembloids from co-culture of cerebral organoids with iMG cells

5 x 10^5^ iHPCs (CD43^+^ cells) were added to medium containing single brain organoids for 15 days promote integration of CD43^+^ cells into the organoid and allow their differentiation into hiMG cells.

### Human Neural Stem Cell (hNSC) culture

Human NSCs derived from H9 hESCs were purchased from Gibco (N7800200) and cultured on Matrigel- or CELLStart-coated plates in knockout D-MEM/F-12 media containing 2 mM GlutaMax, 20 ng/ml bFGF, 20 ng/ml EGF, and 2% StemPro Neural Supplement.

### Neuronal differentiation

To differentiate hNSCs into neurons, hNSCs were plated at 2.5 –5 × 10^4^ cells/cm^2^ in polyornithine- and laminin-coated culture dishes in complete StemPro NSC SFM. After 2 days, the media was changed to neural differentiation medium consisting of neurobasal medium supplemented with B-27 Serum-Free Supplement (1X), GlutaMAX-I (1X) and antibiotic -antimycotic (1X) solutions. Media was subsequently changed every 3–4 days for two weeks.

### HIV-1_BaL_ virus preparation and hiMG cell infection

HIV-1_BaL_ plasmid (consisting of the backbone of the HXB-3 strain of HIV-1 IIIB) was obtained through the NIH AIDS Research and Reference Reagent Program (NIH AIDS Reagent # 11414). HIV-1_BaL_ provirus was produced by transfecting 293FT cells (6×10^6^) with 6μg provirus expressing plasmid pBaL using Lipofectamine 2000 (Invitrogen). Supernatants containing virus were harvested 48h later, centrifuged at 1000 x g for 10 min, filtered through 0.45μm filters and stored at -80^0^C. HIV-1_BaL_ particles in supernatants were quantified by ELISA using HIV-1 p24 antigen, as described ^52,81,82^. hESC-derived iMG cells (0.5×10^6^) were infected with HIV-1_Bal_ using an inoculum of 20-50 ng/ml HIV-1 p24 and incubated overnight at 37^0^C. The inoculum was then removed and residual virus was washed away using fresh culture medium. Culture supernatant samples were collected every three days and the medium changed twice weekly till day 12 for ELISA analysis to monitor viral replication (p24) and cytokine release (IL-6) based on the manufacturer’s instructions.

### Immunocytochemistry and Immunofluorescence

hiMG differentiation was assessed by immunocytochemistry using microglia-specific markers. In other analyses, HIV-1_BaL_-infection of hiMG was confirmed by monitoring p24 co-localization with microglial markers. To do so, cells were washed three times in 1X DPBS, fixed in cold 4% PFA for 20 min at room temperature, and washed three times with PBS. Cells were then blocked in blocking solution (1X PBS, 3% BSA, 0.1% Triton X-100) for 1 hr at room temperature followed by overnight incubation at 4^0^C with appropriate primary antibodies diluted in blocking solution. The next day, cells were washed 3 times with PBS for 5 min each and stained with Alexa Fluor-conjugated secondary antibodies at 1:400 for 1 hr at room temperature in the dark. Cells were washed 3 times with PBS and covered with coverslips using DAPI-containing mounting media (Vectashield mounting medium with DAPI).

Assembloids were sectioned and stained by first washing with PBS and then incubation 30 min in cell recovery solution to dissolve surrounding Matrigel. Assembloids were washed again with PBS, fixed 1 hr in 4% paraformaldehyde,washed three times with PBS and incubated in 30% sucrose overnight. The sucrose solution was then removed, and assembloids were washed with PBS and embedded in OCT for sectioning at 20 μm thickness using a cryostat. Mounted slides were stored at 20^0^C until further use. For staining, sections were rehydrated and washed 3 times with room temperature PBS for 5 min each. Heat-mediated antigen retrieval was performed by using citrate buffer (10mM Citrate, 0.05% Tween 20, pH = 6.0) at 70^0^C for 20 min and then samples were allowed to cool to room temperature. After antigen retrieval, slides were washed three times with PBS and then cryosections were blocked 1 hr in 5% BSA in PBS, washed three times with PBS+0.1% Triton X-100, and incubated with primary antibody at 4°C overnight. Sections were then washed three times with PBST to remove primary antibody and then incubated 1 hr incubation with secondary antibody. Sections were then washed three more times to remove secondary antibodies before mounting in Vectashield mounting medium with DAPI following the manufacturer’s instructions. Dilutions for primary antibodies were: mouse anti-GFAP (1:500; Sigma); TREM-2, goat anti-Iba1 (1:100; Abcam ab5076); rabbit anti-Iba1 (1:500, Synaptic System); rabbit anti-human TMEM-119 (1:100; Abcam, ab185333); chicken anti-MAP2 (1:10000, Abcam); mouse anti-β-tubulin III (1:500, Sigma); and mouse anti-P2RY12 (1:200, Sigma)

## ELISA

Enzyme-linked immunosorbent assay (ELISA) was performed using human IL-6 and HIV-1 p24 ELISA kits (R and D) following the manufacturer’s instructions. In brief, 96-well ELISA plates were incubated with IL-6 capture antibody overnight in coating buffer; for the HIV-1 p24, plates were pre-coated with p24 capture antibody. Wells were washed with PBST 5 times and blocked 1 hr with Assay Diluent at room temperature. After 5 more PBST washes, supernatants from METH-treated, HIV-1_BaL_-infected hiMGs or assembloids were collected at 1, 3, 6, 9 and 12 days, and IL-6 standards were added to wells and incubated either overnight at 4°C or 2h at room temperature. Subsequently, wells were washed 5 times with PBST and incubated 1 hr with diluted detection antibody at room temperature. Wells were again washed 5 times before diluted Avidin-HRP was added for 30 min at room temperature. After 5 washes, substrate solution was added for 15 min, followed by stop solution. ELISA plates were read using a Synergy 2 plate reader (Biotek) at 450 nm and 570 nm.

## AUTHOR CONTRIBUTIONS

SKT designed and performed experiments, analyzed the data, and wrote the manuscript draft; GB, ST, and SJ performed experiments; NGC experimental design and manuscript writing; TMR, conceived the overall project and experimental design, and participated in data analyses, interpretations, and manuscript writing.

## CONFLICT OF INTERESTS

All authors have no competing interests to declare.

## ACKNOWLEDGEMENTS

We thank Drs. F. Furnari for the iPS cell lines; Jonathan Karn, Kelly L Jordan-Sciutto and Marcus Kaul for reagents and helpful advice on experiments, and members of the Rana lab for helpful discussions and advice. We thank the NIH AIDS Reagent Program for the HIV_BAL_ construct. This work was supported in part by grants from the National Institutes of Health (DA049524, DA053630, DA058402).

## SUPPLEMENTARY FIGURES

**Figure S1:**
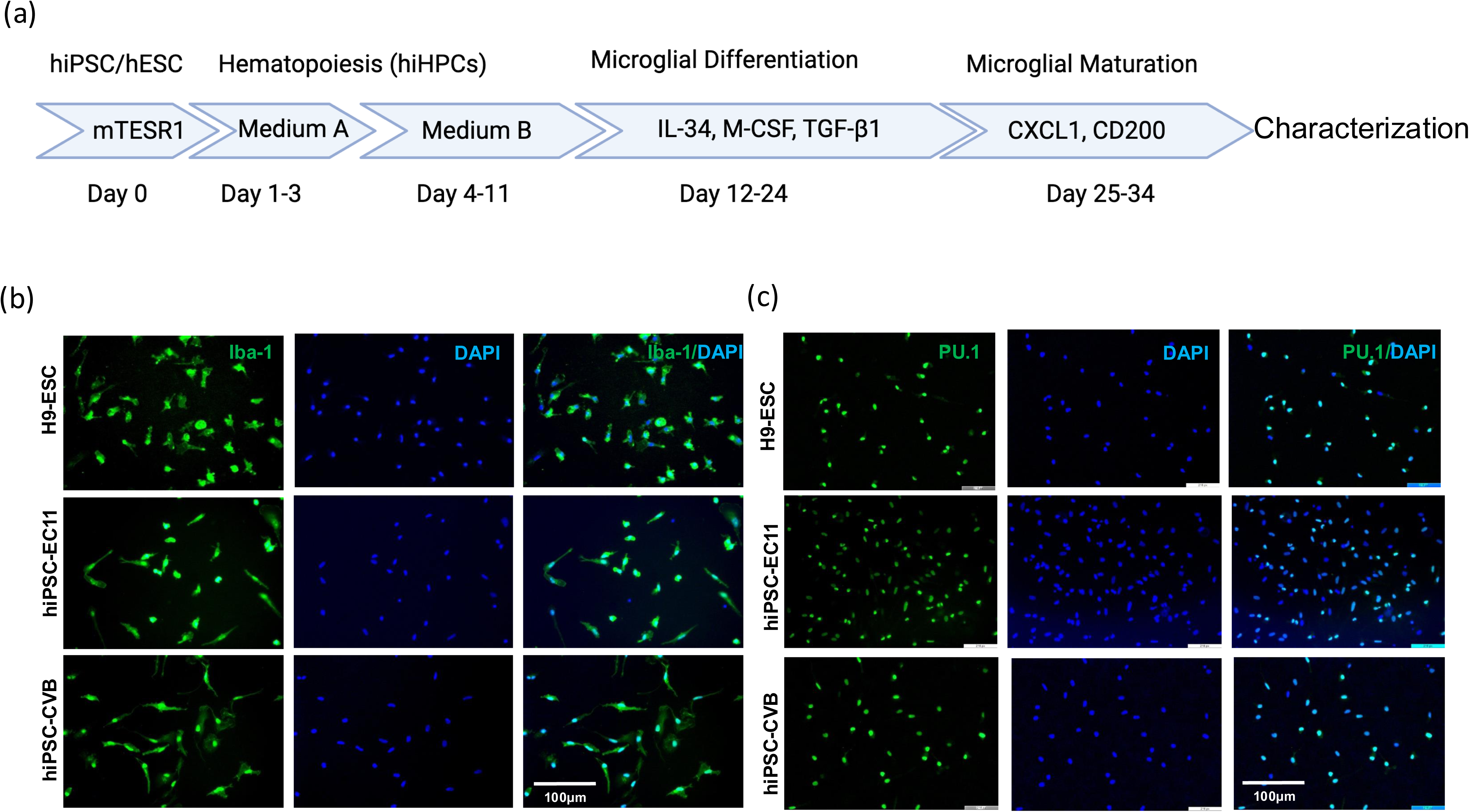
Differentiation and characterization of iMG cells from hESCs/hiPSCs. (a) Representative steps in differentiation of iMG cells as described in methods. (b) (b,c) Immunocytochemical analysis of iMG differentiation from three different stem cell lines (hESC-H9, iPSC-EC11 and iPSC-CVB) showing expression the microglial marker Iba-1 (b) and PU.1 (c), both in green. Samples are also stained with the nuclear marker DAPI (blue). N=3. Scale bar: 100μm.

**Figure S2:**
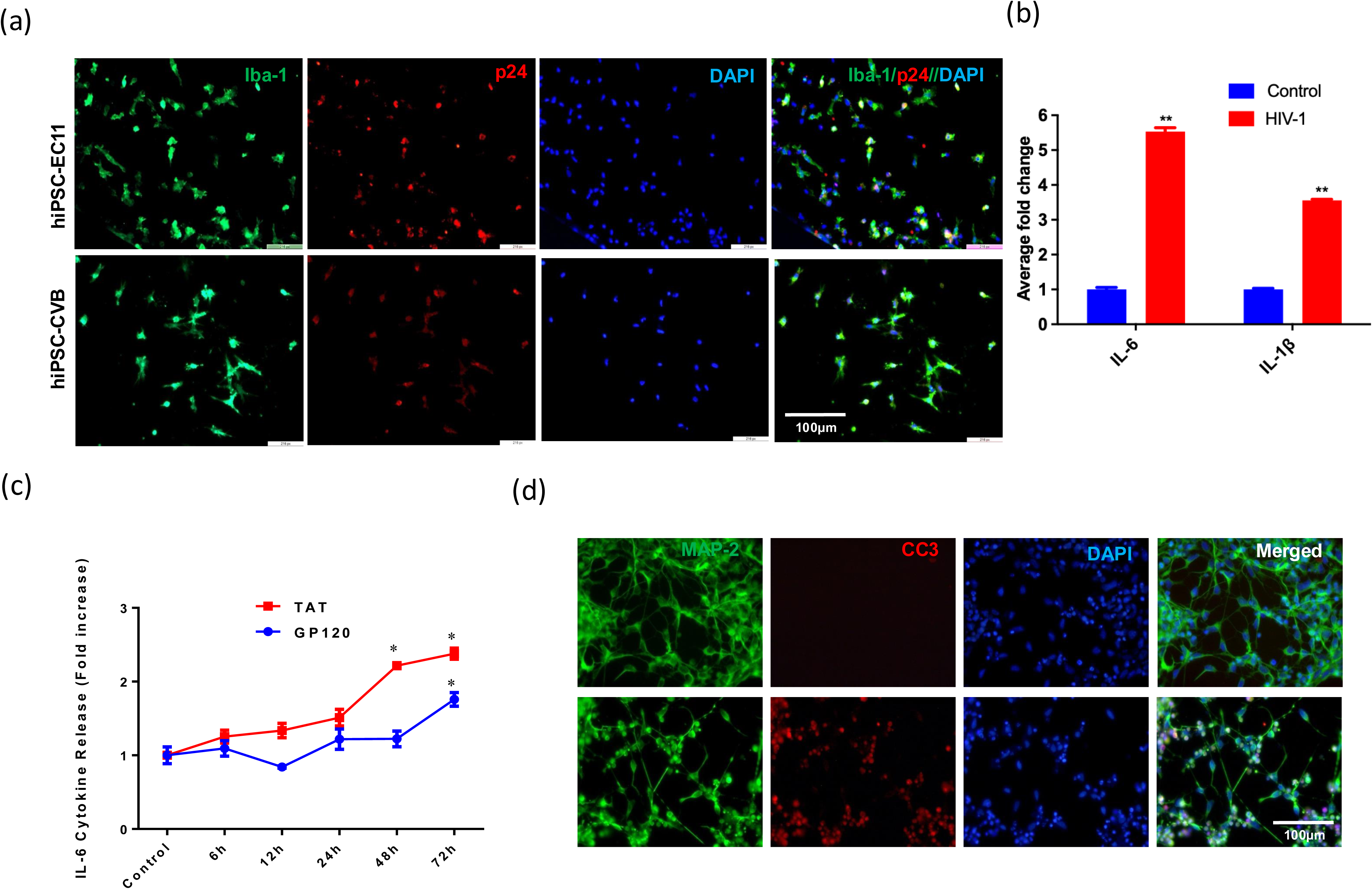
HIV-1 infection of hESC/hiPSC-derived iMG cells and characterization. (a) Immunocytochemical analysis of iMG cells derived the indicated stem cells stained for the microglial marker Iba-1 (green) and the HIV marker p24 (red). Scale bar: 100μm. (b) qRT-PCR analysis of control and HIV-1-infected iMG cells for expression of the pro-inflammatory cytokines IL-6 and IL-1β. **p<0.001, n=3. Error bars indicate mean±SEM. (c) Time-course of IL-6 secretion from iMG cells after treatment with either TAT or GP120, both HIV-1 proteins, as analyzed by human IL-6 ELISA of culture supernatants. n=3, *p<0.01 (d) Immunofluorescence analysis of neuronal cells treated with HIV-1 infected iMG derived culture supernatant and stained for the neuronal marker MAP-2 (green), the apoptosis marker cleaved-caspase-3 (red, CC3), and DAPI. Data suggests that HIV-1-infection may enhance neuronal apoptosis. N=3. Scale bar: 100μm

**Figure S3:**
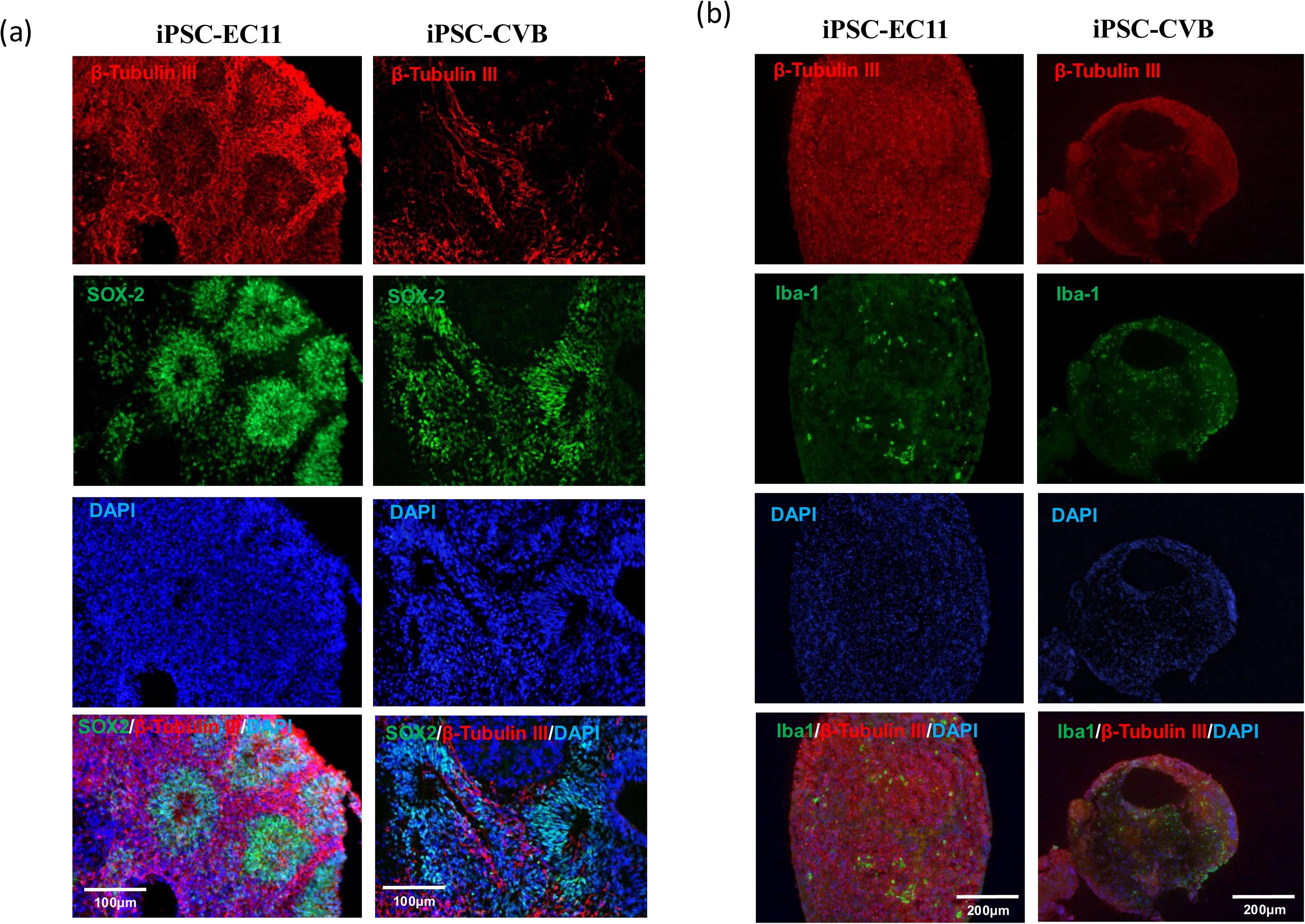
Characterization of hiPSC-derived cortical organoids and 3D cortical assembloids. (a) Immunohistochemical analysis of cortical organoids derived from two different iPSC lines (EC11 and CVB) showing staining for the neuronal marker β-Tubulin III (red), the neural progenitor marker SOX2 (green) and DAPI (blue). n=3. Scale bar: 100μm. (b) Immunohistochemical analysis of samples described in (a) stained for the neuronal marker β-Tubulin III (red), the microglial marker Iba-1 (green) and DAPI. n=3, Scale bar: 100μm.

**Figure S4:**
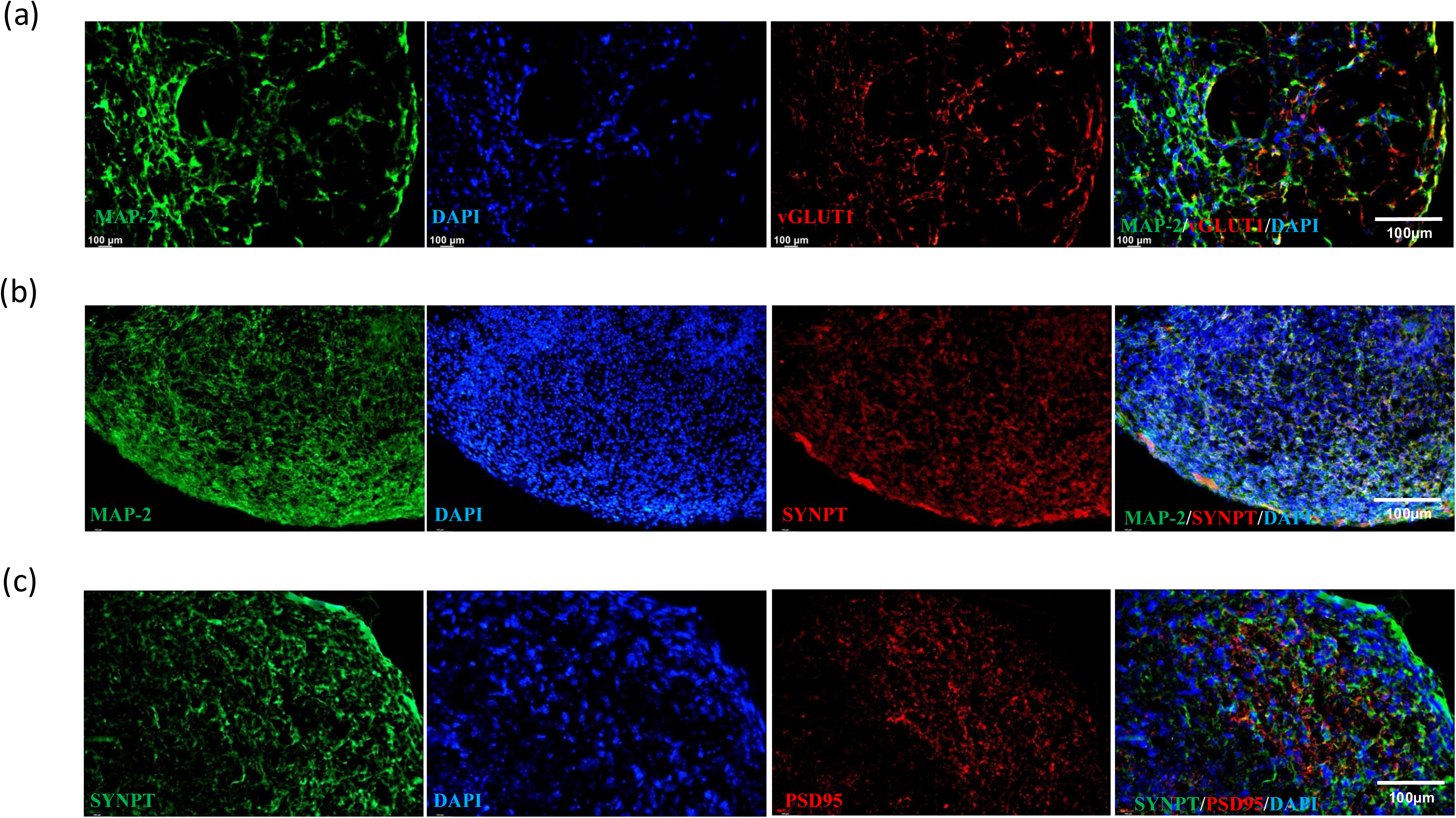
Expression of neuronal markers in hiPSC-derived 3D cortical assembloids. (a) Representative immunohistochemical images of sections from assembloids stained for the neuronal marker MAP2 (green), the glutamatergic neuron marker vGLUT1 (red), and DAPI (blue). n=3. Scale bar: 100μm. (b) Immunohistochemical analysis of tissue sections described in (a) stained for the neuronal marker MAP2 (green), the synaptic marker synaptophysin (SYNPT, red), and DAPI (blue). n=3. Scale bar: 100μm. (c) Immunohistochemical analysis of tissue sections described in (a) stained for SYNPT, (green), the post-synaptic density protein PSD95 (red), and DAPI (blue). n=3, Scale bar: 100μm.

## Notes

### Competing Interest Statement

The authors have declared no competing interest.

